# Cetacean habitat modelling to inform conservation management, marine spatial planning, and as a basis for anthropogenic threat mitigation in Indonesia

**DOI:** 10.1101/2020.07.14.203240

**Authors:** Achmad Sahri, Mochamad Iqbal Herwata Putra, Putu Liza Kusuma Mustika, Danielle Kreb, Albertinka J. Murk

## Abstract

Indonesia harbours a high diversity of cetaceans, yet effective conservation is hampered by a lack of knowledge about cetacean spatial distribution and habitat preferences. This study aims to address this knowledge gap at an adequate resolution to support national cetacean conservation and management planning. Maximum Entropy (Maxent) modelling was used to map the distribution of 15 selected cetacean species in seven areas within Indonesian waters using recent cetacean presence datasets as well as environmental predictors (topographic and oceanographic variables). We then combined the individual species suitable habitat maps and overlaid them with provincial marine spatial planning (MSP) jurisdictions, marine protected areas (MPAs), oil and gas contract areas, and marine traffic density. Our results reflect a great heterogeneity in distribution among species and within species among different locations. This heterogeneity reflects an interrelated influence of topographic variables and oceanographic processes on the distribution of cetacean species. Bathymetry, distance to-coast and −200m isobaths, and Chl and SST were important variables influencing distribution of most species in many regions. Areas rich in species were mainly related to high coastal or insular-reef complexity, representing high productivity and upwelling-modified waters. Although some important suitable habitats currently fall within MPAs, other areas are not and overlap with oil and gas exploration activities and marine traffic, indicating potentially high risk areas for cetaceans. The results of this study can support national cetacean conservation and management planning, and be used to reduce or avoid adverse anthropogenic threats. We advise considering currently unprotected suitable cetacean habitats in MPA and MSP development.

## 1. Introduction

Biodiversity conservation and area-based management call for adequate-resolution species distribution data. Understanding the spatial distribution and habitat preferences of marine mammal species is a high priority for effective conservation management. A principal phase underpinning marine spatial planning (MSP) involves mapping the spatial distribution of ecological processes and biological features (Ehler and Douvere, 2007; Metcalfe et al., 2018). Protecting important habitats of top marine species is also a priority issue for MSP development (Hooker et al., 2011). However, often this information is lacking, inhibiting conservation efforts of these marine species. A sound management plan which adopts the most effective measures to protect marine mammals is clearly constrained by the knowledge on the species’ critical habitats.

Cetaceans are well-known as charismatic species, and as top predators they could have strong effects on community structure and function (Foley et al., 2010). The management of protected areas designed for top predators as umbrella species is highly efficient, resulting in higher biodiversity and more ecosystem benefits (Sergio et al., 2008). However, there is a long history of anthropogenic impacts on cetaceans through whaling, habitat degradation and increased activities in the marine environment that could adversely affect cetaceans (Weir and Pierce, 2013). Cetaceans are particularly susceptible to human threats because of their life-history traits i.e. late maturity and low reproductive rate (Passadore et al., 2018). The variety of anthropogenic pressures includes interactions with fisheries (entanglement, bycatch, prey depletion) (Read, 2008; Reeves et al., 2013), physical and acoustic disturbance mainly by marine traffic (ship strikes, underwater noise) (Erbe et al., 2019; Pennino et al., 2017), seismic activities from oil and gas exploration and naval sonars (Henderson et al., 2014; Rosenbaum and Collins, 2006), and pollution (oil spills, plastic debris, heavy metals, and other chemicals) (Allen et al., 2011; Monk et al., 2014; Tanabe, 2002; Venn-Watson et al., 2015). New threats have also been recognized, including coastal-offshore development and energy production, resource extraction, tourism, and climate change (MacLeod, 2009; Passadore et al., 2018). A prominent stressor to cetaceans is marine traffic which is to some extent linked to shipping (Coomber et al., 2016). Large whales are vulnerable to collisions with vessels throughout the world’s oceans (Laist et al., 2001). In addition, marine activity due to oil and gas exploration and extraction is also growing (Dawson et al., 2018) and this has a major impact on marine life and environment (Azzellino et al., 2012), mainly through noise and potential pollutants. Understanding how both marine traffic and oil-gas concessions overlay with marine mammals’ distributions is crucial for improving future decision-making regarding the zoning of multiple-use MPAs and MSPs.

The Indonesian archipelago holds a high diversity of cetacean species with 34 cetacean species recorded so far (Mustika et al., 2015), accounting for more than a third of cetacean species worldwide (Jefferson et al., 2015). The archipelago was previously one of the largest global whaling grounds (Townsend, 1935) and is highly biologically productive. Indonesia is situated in an upwelling system, where wind regimes and oceanic currents strongly influence the temperature and primary productivity (Drushka et al., 2010; Steinke et al., 2014), benefiting marine species at all levels of the food web, including cetaceans (Huffard et al., 2012). Knowledge on cetacean distribution in Indonesia, however, is not homogeneous, spatially or temporally. There have been relatively few recent cetacean surveys in Indonesia due to its vast area, remote offshore locations, poor weather and sea conditions, limited financial resources for research, and other logistic constraints (Evans and Hammond, 2004). Several locations have received survey coverage, mainly in Lesser Sunda (including Bali and Solor-Alor), Papua, East Kalimantan, West Sumatra and Banda Sea (Ender et al., 2014; Kreb et al., 2015; Kreb and Budiono, 2005; Mustika, 2006; Sahri et al., 2014). These surveys were often undertaken within the framework of wildlife conservation through the establishment of MPAs. No studies have yet assessed the distribution of cetaceans across the whole of the Indonesian archipelago.

All marine mammal species in Indonesia are under strict protection according to national regulations (Sahri et al., 2020), where all species are listed in the annex of the Government Regulation No. 7/1999 (The Government of The Republic of Indonesia, 1999). Based on the online database of the International Union for Conservation of Nature (IUCN, 2020), more than a fifth of the cetacean species that occur in Indonesian waters are listed as ‘threatened’. Information about the spatial distribution of these species is still very limited, yet determining areas that require protection at appropriate scales for national management requires a finer understanding of the species habitat (Dransfield et al., 2014; Redfern et al., 2006). Typically, however, such information is only available in broad geographic regions and usually only in a coarse resolution and for a limited number of species (Kaschner et al., 2016, 2006). In Indonesia, no distribution maps are available at eco-regional or seascape scales. The need for mapping critical habitat for cetaceans in Indonesia was indicated by the National Plan of Action (NPOA) for Cetaceans (Mustika et al., 2015) and under the Regional Plan of Action (RPOA) for the Coral Triangle Initiative (CTI-CFF, 2009) as well as national legislation (MMAF, 2018). They ask for a better understanding of the biology and ecology of cetaceans for conservation purposes. Knowledge on distribution and habitat preference would enable stakeholders to minimize harmful human and cetacean interactions and implement spatially explicit conservation measures in a certain region. Combining cetacean habitat maps and maps of human activities to identify areas of high impact has not been attempted for Indonesia before.

Species distribution models (SDMs) are increasingly used in conservation planning and wildlife management, including for cetaceans (Hammond et al., 2013), especially in developing MSP and designing MPAs (Cañadas et al., 2002) and identifying areas of potential conflict between human activities and marine organisms (Guisan et al., 2013). The models can provide a finer spatial resolution than traditional abundance estimates (Becker et al., 2012). One of the powerful modelling tools used by numerous studies for cetacean species worldwide is Maximum Entropy (Maxent) (Breen et al., 2016). As a presence-only technique, Maxent is particularly useful for studies of species with large ranges and small sample sizes, for regions where systematic surveys are sparse and/or limited in coverage, and for datasets for which absence or effort data are not available (Elith et al., 2011; Moura et al., 2012). Using this modelling technique, it is possible to predict suitable habitats for a range of species and ultimately map areas of high cetacean diversity that is particularly useful for managers and decision-makers (Becker et al., 2012).

This study aims to provide adequate-resolution maps of cetacean distribution and habitat suitability in Indonesia and assess areal overlaps with MPAs, MSPs, and two anthropogenic threats i.e. oil and gas concessions and marine traffic. The information from this study is crucial, and is particularly needed by the Indonesian government to be able to manage their more national scale habitats to ensure species protection as well as to guide policies mitigating anthropogenic threats.

## 2. Materials and Methods

### 2.1. Study area

In this study seven regions within Indonesian waters are included: Sunda Strait (SS), Balikpapan Bay (BB), NE Borneo Seascape (NEBS), SE Sulawesi Seascape (SESS), Lesser Sunda Ecoregion (LSE), Bird’s Head Seascape (BHS), and Fakfak Seascape (FS) (Figure 1). These regions are chosen based on sufficient sighting data availability. The boundaries of the regions used either ecoregions or seascapes depending on the geographical distribution and the extent of presence data, with the exception of the first two regions that were determined based on the MSP jurisdiction (∼12 nm from coastlines). Ecoregions or seascapes were chosen since their boundaries are scientifically-determined and ecologically based (Green and Mous, 2008).

**Figure 1.**
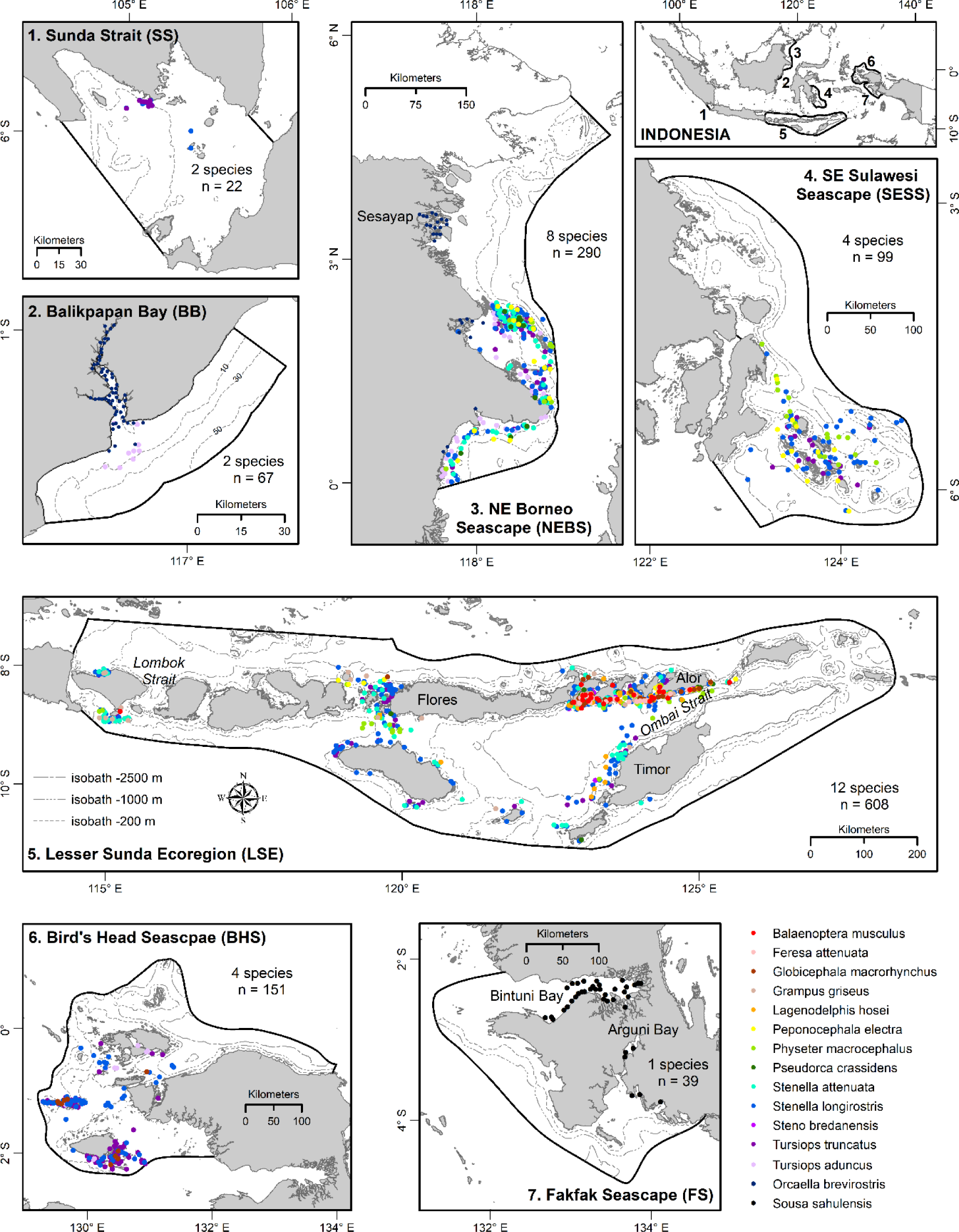
Seven regions in Indonesia that were chosen as the study area with the reported presences (total n=1276) of the 15 selected cetacean species used in Maxent. Isobaths (−200, −1000, and −2500m) as indicated in the Lesser Sunda Ecoregion map legend also apply for other regions, except for Balikpapan Bay which has its own isobaths. Number of species and total number of reported presence points are given in each region map.

### 2.2. Sighting data

Species presence data was collected from multiple sources from several cetacean programs in Indonesia from 2000-2018 (Figure 1). These are currently the best sources of information on the occurrence of cetaceans in the Indonesian archipelago. The list of species, the number of sightings, and the number of sightings used in SDMs, however, were different in each region (Table S1). The data was collected from both dedicated surveys and non-systematic surveys from a range of platforms of opportunity. In order to improve the number of sightings, records from several surveys were pooled, regardless of the survey methods. Long term and pooled data capture both inter-seasonal and inter-annual variability in cetacean distribution (Cotté et al., 2010). Only species with >10 sightings in a region were included in SDMs, as this is the minimum number of sightings for which the outset constant accuracy of SDMs can be reached (Wisz et al., 2008). This treatment resulted in 15 species out of 34 being used for this study (Table S1).

To avoid model over-fit, occurrence data should be spatially independent (free from spatial auto-correlation). An over-fitted model will reduce the model’s ability to predict spatially independent data and inflate model performance values (Boria et al., 2014). The ‘Spatially rarefy occurrence data tool’ in SDMtoolbox 2.0 (Brown et al., 2017) was used to spatially filter locality data by a certain distance, reducing occurrence localities to a single point within the specified Euclidean distance or according to spatial heterogeneity of environmental variables. The maximum distance of 3 km (1 km for two smaller regions: the SS and BB) was used for sighting rarefying because of the high spatial heterogeneity in this study area and to prevent loss of extensive occurrence data. This graduated filtering method is particularly useful for this study, since most sighting records were concentrated in certain locations as a consequence of non-systematic surveys, so they might be spatially correlated. After the rarefying process, a total of 1276 presence-only data points of the original 2252 data for 15 cetacean species were used in Maxent.

### 2.3. Environmental predictors

Ten variables are available in the study area (Table 1) that can be categorised into two types i.e. topographic variables (bathymetry, slope, distance- to coast, shelf, −200m, −1000m, and −2500m isobaths), and oceanographic variables (sea surface temperature (SST), sea surface salinity (SSS), chlorophyll-a concentration (Chl)). These environmental features have been used as proxies to understand cetacean habitat in many studies (Azzellino et al., 2012; Fiedler et al., 2018; Tardin et al., 2019; Viddi et al., 2010). The collinearity among variables was checked in each region and only variables with Pearson’s correlation values less than 0.75 were included in ecological modelling. The ‘Remove highly correlated variables tool’ in SDMtoolbox 2.0 was used to check the collinearity among variables and eliminate correlated variables. Therefore, the selected variables used in modelling were different in each region (Table 2).

**Table 1.** Environmental predictors available for the study areas and the information sources

| Predictors | Unit | Source |
| --- | --- | --- |
| Bathymetry | m | GEBCO (https://www.gebco.net) |
| Slope | % | GEBCO and ArcGIS derived |
| Distance to isobaths: |  |  |
| -200 m (d_200) | km | GEBCO and ArcGIS derived |
| -1000 m (d_1000) | km | GEBCO and ArcGIS derived |
| -2500 m (d_2500) | km | GEBCO and ArcGIS derived |
| Distance to: |  |  |
| coast (d_coast) | km | Indonesian Geospatial Information Agency and ArcGIS derived |
| shelf (d_shelf) | km | Seafloor Geomorphic Features Map (Harris et al., 2014) and ArcGIS derived |
| Oceanographic predictors: |  |  |
| Sea surface temperature (SST) | °C | Aqua MODIS (https://oceancolor.gsfc.nasa.gov) |
| Sea surface salinity (SSS) | PSU | SODA 3.3.1 (http://apdrc.soest.hawaii.edu) |
| Chlorophyll-a (Chl) | mg.m<sup>-3</sup> | Aqua MODIS (https://oceancolor.gsfc.nasa.gov) |

**Table 2.** Environmental predictors used per area in the Maxent models (unless removed based on multi-collinearity test, indicated with ‘*’)

| Predictors | SS | BB | NEBS | SESS | LSE | BHS | FS |
| --- | --- | --- | --- | --- | --- | --- | --- |
| Bathymetry | v | v | v | v | v | v | v |
| Slope | v | v | v | v | v | v | v |
| Distance to isobaths: |  |  |  |  |  |  |  |
| -200 m (d_200) | v | n.a | v | v | v | v | * |
| -1000 m (d_1000) | v | n.a | v | v | v | n.a | n.a |
| -2500 m (d_2500) | n.a | n.a | * | v | v | v | n.a |
| Distance to: |  |  |  |  |  |  |  |
| coast (d_coast) | v | n.a | v | * | * | v | v |
| shelf (d_shelf) | * | n.a | * | * | * | * | * |
| Oceanographic predictors: |  |  |  |  |  |  |  |
| Sea surface temperature (SST) | v | v | * | v | v | v | v |
| Sea surface salinity (SSS) | * | v | v | v | v | * | v |
| Chlorophyll-a (Chl) | v | * | v | v | v | v | * |

n.a : predictor was not available for the specific region, so was not used in the modelling
'v' indicates the variables that were used in the models in each region
SS: Sunda Strait, BB: Balikpapan Bay, NEBS: NE Borneo Seascape, SESS: SE Sulawesi Seascape, LSE: Lesser Sunda Ecoregion, BHS: Bird's Head Seascape, FS: Fakfak Seascape.

Bathymetry data with a 1 km^2^ grid was obtained from the General Bathymetric Chart of the Oceans (GEBCO, https://www.gebco.net). Slope was derived from the GEBCO using the ‘Spatial Analyst extension’ in ArcGIS 10.3 (Environmental Systems Research Institute, Inc.). The coastlines were obtained from the Indonesian Geospatial Information Agency, while submarine shelf was acquired from the Seafloor Geomorphic Features Map (Harris et al., 2014). Isobaths of −200m, −1000m, and −2500m were generated from the bathymetry data using the ‘Contour tool-Spatial Analyst extension’ in ArcGIS 10.3. Distance to-coast, shelf, and the three isobaths were generated using the ‘Euclidean Distance tool-Spatial Analyst extension’ in ArcGIS 10.3. The SST and Chl data were downloaded from Aqua MODIS (https://oceancolor.gsfc.nasa.gov), while the SSS data was downloaded from SODA 3.3.1 (http://apdrc.soest.hawaii.edu). Mean annual SST, Chl, and SSS from daily records covering the same time frame as the sightings data were used. Not enough sightings distributed over the seasons were available for some species to average monthly or seasonal differences. The ‘Inverse distance weighted (IDW) algorithm’ in ArcGIS 10.3 was used to interpolate the spatial data of these oceanographic variables. The selection of spatial resolutions for final environmental variables was primarily based on data availability. Since bathymetry and slope were already in a 1 km^2^ grid, the other variables were aggregated to match the same grid size and cover the same area in each region.

### 2.4. Maxent model setting

Maxent software version 3.4.1 (https://biodiversityinformatics.amnh.org/open_source/maxent) was used to generate probabilistic predictions and habitat models for each cetacean species. This software is a maximum entropy algorithm, specifically developed for presence-only data (Phillips et al., 2006). Maxent has been successfully applied in situations where absence data was not available (Elith et al., 2011, 2006), and was widely used when working with combined data collected with different methodologies, as done in the present study. Maxent estimates the relative probability distribution of species occurrence by finding the probability distribution of maximum entropy, i.e. the distribution that is closest to uniform across the study area. The probability of occurrence can be interpreted as an estimate of the probability of the presence under a similar level of sampling effort as used to obtain the known occurrence data (Phillips and Dudík, 2008).

The following Maxent settings were chosen with regard to data limitations and the specific questions of the study (Merow et al., 2013): i) logistic output to easily understand where the model predicts the occurrence of each cetacean species; ii) 30 % random test percentage; iii) default regularization parameters, auto feature class types, and 500 maximum iterations; iv) 10-fold bootstrap replicated run type, a setting that allows replacement in sampling replicates and is particularly useful when the number of sightings are low (Fielding and Bell, 1997); and v) the maximum number of background points was 10,000 (over 43,800-369,861 available points) as number of background points greater than 10,000 does not improve the predictive ability of the model (Phillips and Dudík, 2008). The maximum number of background points for two smaller regions (the SS and BB) was set to 1,720 and 320 respectively (∼13% of its own available points, comparable to the average ratio of background-available points of other regions). Bias files were used to refine background point selection in Maxent. Bias files constrain the location and density of background point sampling to ensure that background points are generated from the same environmental space as the presence locations and allow the user to account for collection sampling bias (Phillips et al., 2009). Bias files were created and hence background selection was carried out using 30 km (15 km for two smaller regions: the SS and BB) buffered local adaptive convex-hull for individual species using the ‘Background selection tool’ in the SDMtoolbox 2.0. The buffers chosen have been shown to best restrict background point selection within the environmental space.

The performance of each Maxent model was evaluated using the AUC (area under the receiver-operating-characteristic curve), which assesses model discriminatory power by comparing model sensitivity (i.e., true positives) against model 1 minus specificity (false positives) from a set of test data (Phillips et al., 2006). The AUC value provides a threshold-independent metric of overall accuracy, and ranges between 0 and 1. An AUC value above 0.5 indicates that the model performs better than random (Phillips and Dudík, 2008), while values between 0.6 and 0.9 were indicative of a well fitted model (Breen et al., 2016). To assess how much each environmental variable contributed to the Maxent run, jackknife tests of variable importance were conducted by running the model with-only and without a variable at a time. It was possible to evaluate the contribution (gain) of each variable with respect to the whole ensemble of variables, and to evaluate the effects of the lack of the selected variable on the model compared to the set of overall variables (Elith et al., 2006). To distinguish suitable and unsuitable habitats, the ‘maximum training sensitivity plus specificity threshold’ was applied to the predicted distribution maps in ArcGIS 10.3. The mean threshold of 10 replicates of each species model was used as a binary threshold for presence/absence of corresponding species, above which a suitable habitat is considered to occur. This is the point where the proportion of correctly predicted presences and absences are maximized (Liu et al., 2005).

Maxent has a common problem with producing over-prediction, since it gives higher probability scores for habitat suitability in areas with similar environment characteristics to the sighting locations, although the predicted areas are located outside the observed range. To overcome the over-prediction, minimum convex polygon (MCP) with 30 km (15 km for two smaller regions: the SS and BB) as a buffer distance from the sighting point of each species was used in SDMtoolbox 2.0. This technique resulted in model outputs that represent the suitable habitat within an area of known occurrence, excluding potentially suitable habitats outside the observed range and unsuitable habitats throughout the study areas.

### 2.5. Species combined maps overlap with spatial conservation management system and anthropogenic threats

Firstly, a combined suitable habitat map (hereafter called ‘combined map’) was made by summing individual binary species suitability maps in each region (Brown et al., 2017). This map, however, does not necessarily represent the absolute species richness (Brown, 2014) as only data from the selected 15 out of the 34 reported species was used. Next, the areal overlap between cetacean habitats and current spatial conservation management system was made by superimposing the combined map in each region with maps of MPAs and MSP jurisdiction. MPA polygons were gained from the Ministry of Marine Affair and Fisheries of Indonesia, while MSP jurisdiction (12 nm from coastlines, (The Government of The Republic of Indonesia, 2014a)) was generated using ‘Buffer-Analysis Tools’ in ArcGIS 10.3. Similarly, to predict cetacean exposure to anthropogenic threats, the combined map in each region was overlaid with oil and gas concession areas and marine traffic. Oil and gas concession areas in Indonesia from Patra Nusa Data (2016) were digitized in ArcGIS 10.3. Marine traffic was indicated by shipping density from a global map of shipping traffic (Halpern et al., 2008).

## 3. Results

### 3.1. Maxent model performance

The Maxent model performed well for the majority of species modelled for most regions. All model outputs presented good discriminant power with AUC scores ranging from 0.740 to 0.898, thus can be considered appropriate models. The standard deviation of the AUCs showed low values less than 0.05 for 29 out of the total 33 model outputs (Figure 2). The threshold values used to distinguish suitable and unsuitable habitats ranged from 0.302 to 0.555 (Figure S1). A comparison of suitable habitat from Maxent model predictions with the distributions of sightings shows a good agreement for most species in most regions (Figure S1). Of the 33 Maxent model outputs, six were performed based on only a few presence data points (between 10 and 15 data points) (Figure 2).

**Figure 2.**
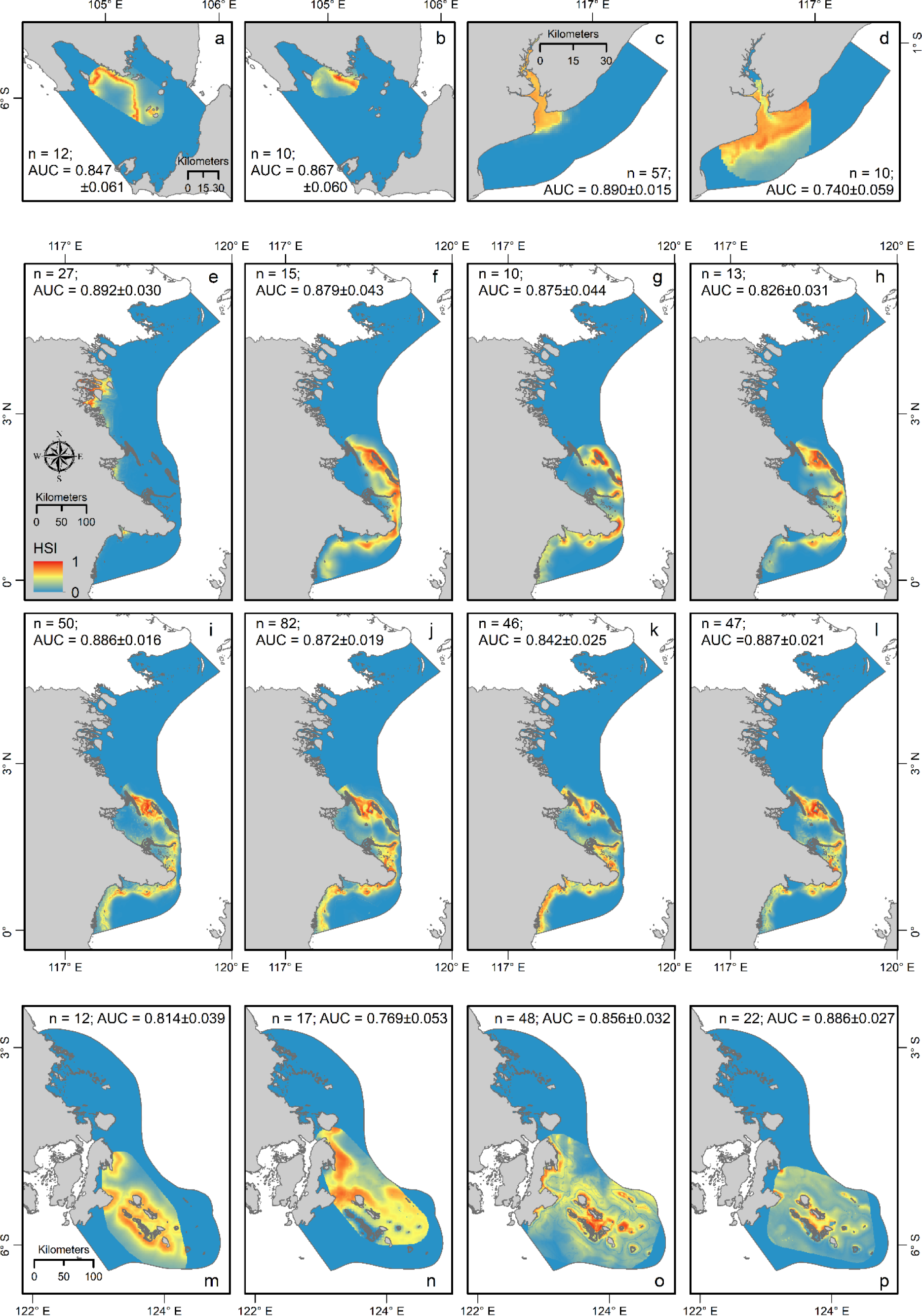
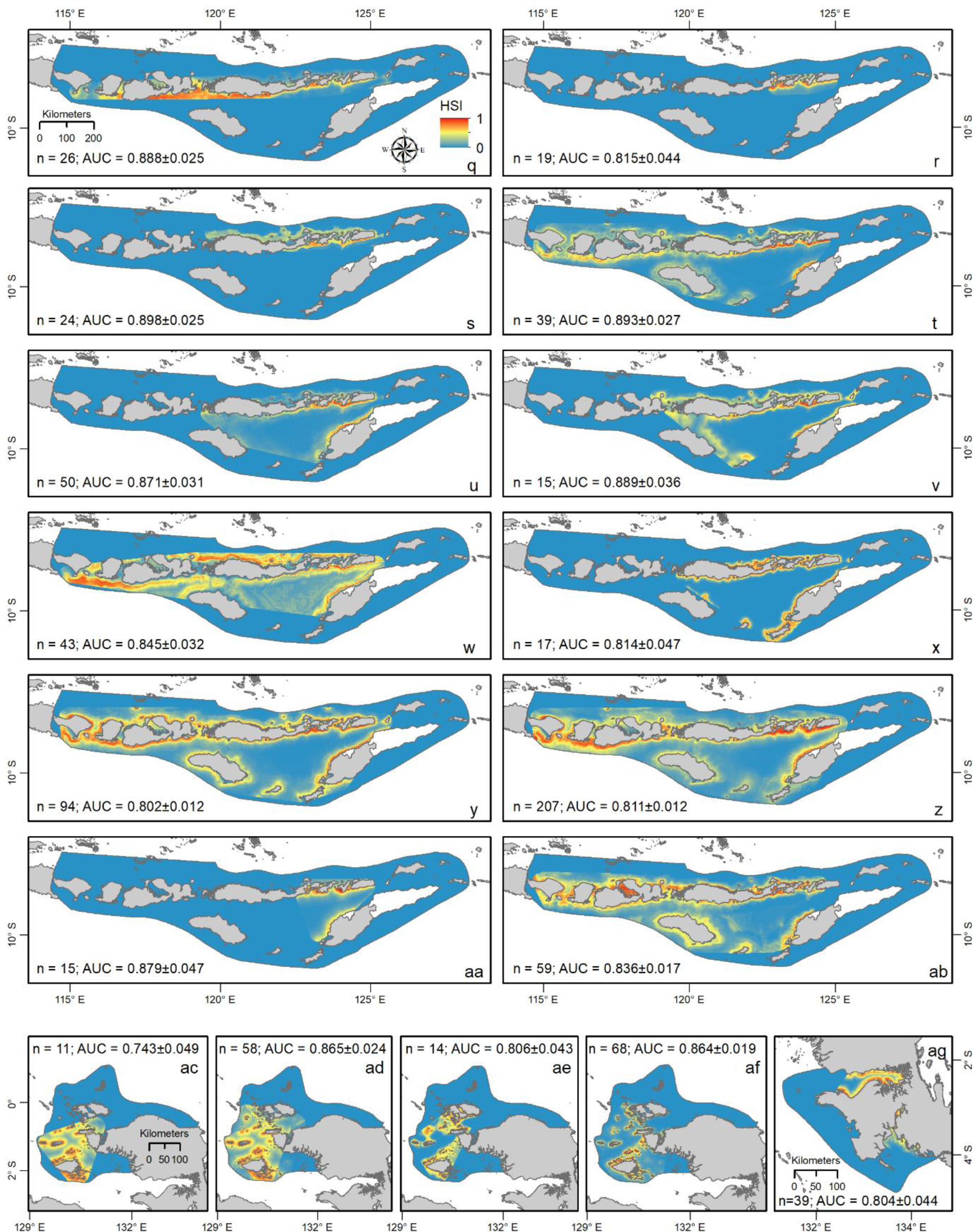
Individual spatial distribution of 15 cetaceans in 7 regions in Indonesia. Habitat Suitability Index (HSI) range from 0-1. Species names are indicated by letters: *Balaenoptera musculus* (q), *Feresa attenuata* (r), *Globicephala macrorhynchus* (s,ac), *Grampus griseus* (t), *Lagenodelphis hosei* (u), *Orcaella brevirostris* (c,e), *Peponocephala electra* (f,m,v), *Physeter macrocephalus* (g,n,w), *Pseudorca crassidens* (h,x), *Sousa sahulensis* (ag), *Stenella attenuata* (i,y), *Stenella longirostris* (a,j,o,z,ad), *Steno bredanensis* (aa), *Tursiops aduncus* (d,k,ae), and *Tursiops truncatus* (b,l,p,ab,af). n = number of sightings, AUC is area under ROC curve.

### 3.2. Predicted distribution, important variables and habitat preferences

Our results reflected a great heterogeneity in distribution among species and within species among different regions (Figure 2). Within the same region, predicted distributions varied between species and some species showed areas of overlap. The predicted distributions showed a similar pattern for overlapping species in a region, although differences in spatial extent were identified between species (Figure 2).

Important variables differed across the model outputs for each species and region (Table 3). In general, bathymetry, distance to coast, distance to −200m isobath, Chl, and SST were important variables for most species in many regions. High predictive scores or habitat suitability index (HSI) of most species (Figure 2) were mainly in areas that indicate the presence of these environmental predictors that represent productive areas. Habitat preference also varied among species (Tables S2-S8). Below, the distributions of cetacean species, the important variables and habitat preferences are elaborated upon per region.

**Table 3.** Relative importance (%) of each environmental variable in explaining the species distribution in the Maxent model. Bold numbers in grey background are the two most important variables, bold italic numbers with grey background are the third or the fourth important variables that were required to achieve a summed explanation of at least 50%. The variable names in the first row are the names given in Table 1.

|  | Bathymetry | Slope | d_200 | d_1000 | d_2500 | d_coast | SST | SSS | Chl |
| --- | --- | --- | --- | --- | --- | --- | --- | --- | --- |
| **Sunda Strait (SS)** |  |  |  |  |  |  |  |  |  |
| *Stenella longirostris* | **13** | 4.8 | **76.2** | 3.2 | - | 0 | 2.7 | - | 0.1 |
| *Tursiops truncatus* | 0.4 | 3.8 | **42.9** | 4.9 | - | 14.5 | 6.9 | - | **26.6** |
| **Balikpapan Bay (BB)** |  |  |  |  |  |  |  |  |  |
| *Orcaella brevirostris* | 7 | 4 | - | - | - | - | **74** | **15** | - |
| *Tursiops aduncus* | 12 | **41.5** | - | - | - | - | 13.3 | **33.3** | - |
| **NE Borneo Seascape (NEBS)** |  |  |  |  |  |  |  |  |  |
| *Orcaella brevirostris* | **13** | 8.5 | 3 | 4.7 | - | **56.6** | - | 10.9 | 3.3 |
| *Peponocephala electra* | 12 | 3.5 | **49.4** | 15.7 | - | 1.2 | - | 1.5 | **16.7** |
| *Physeter macrocephalus* | 0.8 | 1.4 | **27.3** | 7.1 | - | **53.1** | - | 10.3 | 0 |
| *Pseudorca crassidens* | 0.9 | **42.6** | 12.6 | 4.3 | - | 16.3 | - | 5 | **18.3** |
| *Stenella attenuata* | **17.2** | 13.3 | 15.3 | **16.3** | - | 11.8 | - | **17.3** | 8.8 |
| *Stenella longirostris* | **32.2** | 7.6 | 5.6 | 14.6 | - | 12.6 | - | 6.4 | **21** |
| *Tursiops aduncus* | **16.2** | 12.8 | 5.5 | 5.4 | - | **47.6** | - | 6.9 | 5.6 |
| *Tursiops truncatus* | **18** | 12.7 | 6.2 | 6.6 | - | **23** | - | 12.1 | **21.4** |
| **SE Sulawesi Seascape (SESS)** |  |  |  |  |  |  |  |  |  |
| *Peponocephala electra* | 7.4 | 5 | **43** | 10.2 | 6.2 | - | **16.7** | 9.5 | 2 |
| *Physeter macrocephalus* | 10.4 | 6.9 | 11.3 | **15.6** | 1.7 | - | 0.5 | 3.2 | **50.4** |
| *Stenella longirostris* | 10.3 | 11.3 | **13.3** | **18** | 7.8 | - | **13.3** | 11.6 | **14.4** |
| *Tursiops truncatus* | **57.8** | 4.2 | 4.8 | 6.4 | 5.2 | - | **7.7** | 7.4 | 6.5 |
| **Lesser Sunda Ecoregion (LSE)** |  |  |  |  |  |  |  |  |  |
| *Balaenoptera musculus* | **45.9** | 5.4 | 4.4 | 4.3 | 2.1 | - | 12.1 | 8.7 | **17.1** |
| *Feresa attenuata* | 15.1 | 2.8 | **33.3** | 11.1 | 3 | - | **17** | 6.3 | 11.4 |
| *Globicephala macrorhynchus* | **25.3** | 6.7 | 12.9 | **16** | 7.6 | - | 7.1 | 5.5 | **18.9** |
| *Grampus griseus* | **18** | 8.9 | **30.7** | 10.1 | 9.4 | - | 5.1 | 5.4 | **12.4** |
| *Lagenodelphis hosei* | **33.9** | 2.5 | **19.1** | 4.9 | 12.6 | - | 1.1 | 11.5 | 14.4 |
| *Peponocephala electra* | **25.9** | 0.6 | 8.5 | 23.1 | 9.9 | - | **26.7** | 1.5 | 3.8 |
| *Physeter macrocephalus* | **13.8** | 9.9 | 6.7 | **25.7** | 12.3 | - | **15.1** | 10.3 | 6.2 |
| *Pseudorca crassidens* | **32.6** | 3.8 | **56.8** | 0 | 0 | - | 5 | 0.6 | 1.2 |
| *Stenella attenuata* | **25.3** | 7.9 | **13.6** | 3.8 | 12.5 | - | 8.8 | 11.4 | **16.7** |
| *Stenella longirostris* | **16.9** | 7.4 | **19.8** | 6.1 | 11.9 | - | 6.4 | **19.9** | 11.6 |
| *Steno bredanensis* | 11.8 | 3 | **39.9** | 11.7 | 0.2 | - | **26.8** | 2.3 | 4.3 |
| *Tursiops truncatus* | **32.1** | 4.2 | 14.5 | 10.2 | 4.1 | - | **18** | 2.9 | 14 |
| **Bird's Head Seascape (BHS)** |  |  |  |  |  |  |  |  |  |
| *Globicephala macrorhynchus* | 5 | 0.2 | 12.9 | - | 0.9 | **29.3** | **50.4** | - | 1.3 |
| *Stenella longirostris* | 8.5 | 8.1 | **12.1** | - | 8.4 | **40.7** | 12 | - | 10.2 |
| *Tursiops aduncus* | **68.3** | 5.1 | 6.9 | - | 1.3 | **16.4** | 1.1 | - | 0.9 |
| *Tursiops truncatus* | **14.3** | 4.4 | 6.8 | - | 4 | **53.4** | 8.6 | - | 8.5 |
| **Fakfak Seascape (FS)** |  |  |  |  |  |  |  |  |  |
| *Sousa sahalensis* | **19.1** | 12.3 | - | - | - | **61.4** | 4.2 | 3 | - |

- : predictor was not used in modelling either due to its unavailability in the region, or was removed due to multi-collinearity

In the <u>Sunda Strait (SS)</u>, both *Stenella longirostris* and *Tursiops truncatus*, were distributed near coastal areas, insular areas and around the −200m isobath, although *S. longirostris* was spread wider (Figure 2a,b). Distance to −200m isobath was prominently determining the distribution of *S. longirostris* and *T. truncatus*. The second most important variable explaining the distribution of both species, however, differed; bathymetry for *S. longirostris*, and Chl for *T. truncatus* (Table 3). The habitat preferred by *S. longirostris* was characterized by being closer to −200m isobath, and at areas with a depth of around −255m, and these characteristics were significantly different in comparison with habitat where the species did not occur (Table S2). Similarly, the habitat preference of *T. truncatus* was related to areas closer to −200m isobath, but having higher Chl (Table S2).

In the <u>Balikpapan Bay (BB)</u>, the high predicted distribution of *Orcaella brevirostris* was principally found in river areas (Figure 2c), while *Tursiops aduncus* was distributed both in estuarine and coastal waters (Figure 2d). The distribution of *O. brevirostris* was mainly determined by SST and SSS, while the distribution of *T. aduncus* was dominantly explained by slope and SSS (Table 3). *O. brevirostris* preferred habitat that was characterised by higher SST and lower SSS (Table S3). *T. aduncus* was likely to occur in areas with steeper sea-bottom slope and lower SSS (Table S3).

In the <u>NE Borneo Seascape (NEBS)</u>, the predicted distributions were almost identical for all species and present in broader areas (Figure 2f-l), except *O. brevirostris* which was restricted to estuarine areas (Figure 2e) like in the BB. Most of the delta areas of the NEBS were predicted to be a suitable habitat for *O. brevirostris*, especially in Sesayap in northern areas (Figure 2e; see Figure 1 for detailed location names). Areas that consistently indicate high habitat suitability for the other species were in coastal and complex insular-reef areas. High suitability predictions also extended towards the southern part of the region along the coastline. Bathymetry, distance to coast and Chl were the most important variables determining the distribution of most species (Table 3). The distribution of two deep diving species, *P. electra* and *P. macrocephalus*, was also explained by distance to −200m isobath. The habitat preferences of all species in the NEBS are detailed in Table S4.

In the <u>SE Sulawesi Seascape (SESS)</u>, the model suggested that *Peponocephala electra* and *Physeter macrocephalus* have a wide distribution (Figure 2m,n), with *P. macrocephalus* tending to avoid shallow reefs. *S. longirostris* occurred closer to the complex insular-reef areas, while *Tursiops truncatus* was occupying both coastal and complex reef areas (Figure 2o,p). Some patches of predicted high suitability for *S. longirostris* and *T. truncatus* occur in the vicinity of oceanic islands. Environmental variables that explain most of the distribution of each species are given in Table 3. The two most important variables for *P. electra* were distance to −200m isobath and SST. Chl and distance to −1000m isobath were the major predictors that determined the distribution of *P. macrocephalus*. The distribution of *T. truncatus* was well-explained by bathymetry and SST. *S. longirostris* distribution was determined by four most important variables i.e. distance to −1000m isobath, Chl, distance to −200m isobath, and SST. The preferred habitat areas of these species can be found in Table S5.

In the <u>Lesser Sunda Ecoregion (LSE)</u>, all species exhibited relatively uniform spatial distributions, concentrated along the coastal areas with the differences in spatial extents (Figure 2q-ab). Areas with relatively low occurrence, so relatively unsuitable for most species, were in deeper oceanic waters. Two species, *P. macrocephalus* and *P. electra* however, were predicted to occur in deeper oceanic waters, although with low predicted values. For most species, two topographic variables, bathymetry and distance to −200m isobath, and two oceanographic variables, SST and Chl, were the most important variables (Table 3). The distribution of two deep diving species, *G. macrorhynchus* and *P. macrocephalus*, was also explained by distance to −1000m isobath (Table 3). The habitat preferences of all species in the LSE can be seen in Table S6.

In the <u>Bird’s Head Seascape (BHS)</u>, the predicted distributions were relatively homogeneous and predicted mixed-species distribution overlap, although there was spatial variability among species (Figure 2ac-af). Areas suggested as favourable habitat for all species were dominated by coastal and complex insular-reef areas. Distance to coast and bathymetry were two major environmental variables that drove the distribution of most species. The habitat preferences of the species occurring in the BHS can be found in Table S7.

Finally, for the <u>Fakfak Seascape (FS)</u>, the model showed *Sousa sahulensis* to principally utilise estuarine areas. Two main estuarine areas with consistently high predicted distribution of *S. sahulensis* were the Bintuni Bay and the Arguni Bay (Figure 2ag; see Figure 1 for detailed location names). Areas further offshore seem to be less suitable for the species. The distribution of *S. sahulensis* were determined by distance to coast and bathymetry (Table 3). The preferred habitat of *S. sahulensis* was closer to coast in areas with depths around −23m (Table S8).

#### Habitat preferences of same species in different regions

Our modelling results show great heterogeneity in within-species habitat preferences among different regions. Nine out of 15 species occurred in more than one region (Figure 2). The distribution of *O. brevirostris* mainly occurred in river and estuarine areas both in the BB and the NEBS. The other species that were present in more than one region did not show consistent habitat preferences. For instance, *P. macrocephalus* was associated with more complex insular-reef areas in the NEBS (Figure 2g), utilised more coastal areas in the SESS but avoided shallow reefs (Figure 2n), and occupied both coastal and deeper waters in the LSE (Figure 2w). The heterogeneity in distribution for other species that occurred in more than one region can be seen in Figure 2.

### 3.3. Combined species maps and overlap with management systems and threats

Maxent results can be reported as probabilities or binary output (i.e. suitable vs. unsuitable habitat). The latter is easier to interpret and has important implications for managers to define areas of interest. The combined suitable habitat map of all species (the combined map) in each region shows distinct patterns, with most used common locations along coastlines and complex insular-reef areas. The individual suitable habitat maps for each species in each region are shown in Figure S1. The hotspots based on the combined maps were identified in each region, and most prevalent in the NEBS, SESS, LSE and BHS (Figure 3 and 4). In the NEBS, SESS, and BHS the hotspots can be found around complex insular-reef areas, while in the LSE two hotspots were identified in coastal areas (Figures 3 and 4).

**Figure 3.**
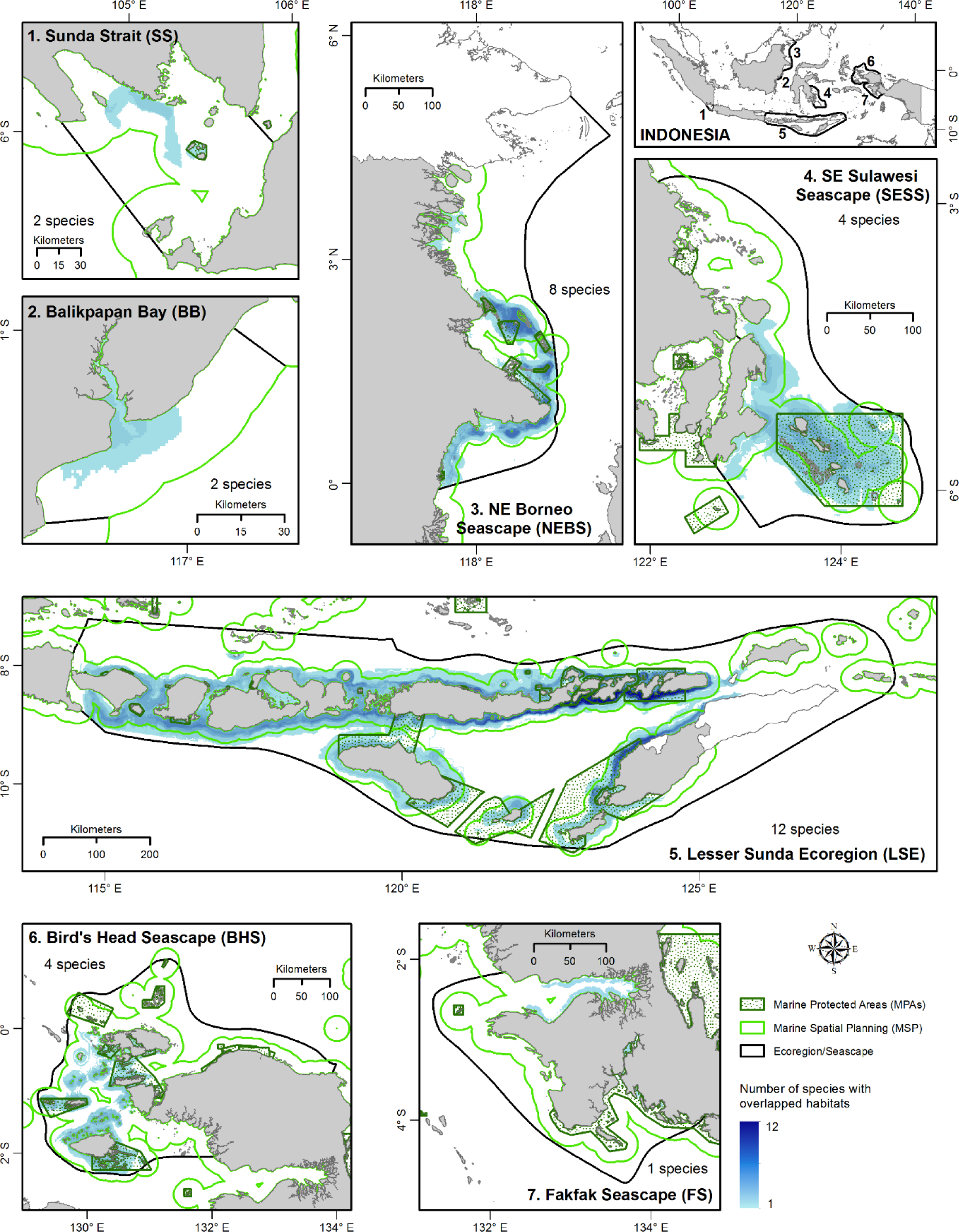
Species combined maps (the suitable habitats for at least one species, gradual blue colours indicating total number of species in each region) and spatial gap analysis with provincial marine spatial planning (MSP) jurisdiction (light green lines), and marine protected areas (dark green dotted areas). The black lines indicate the modelled area (ecoregion or seascape). The number of overlapped species habitats are different in each region.

**Figure 4.**
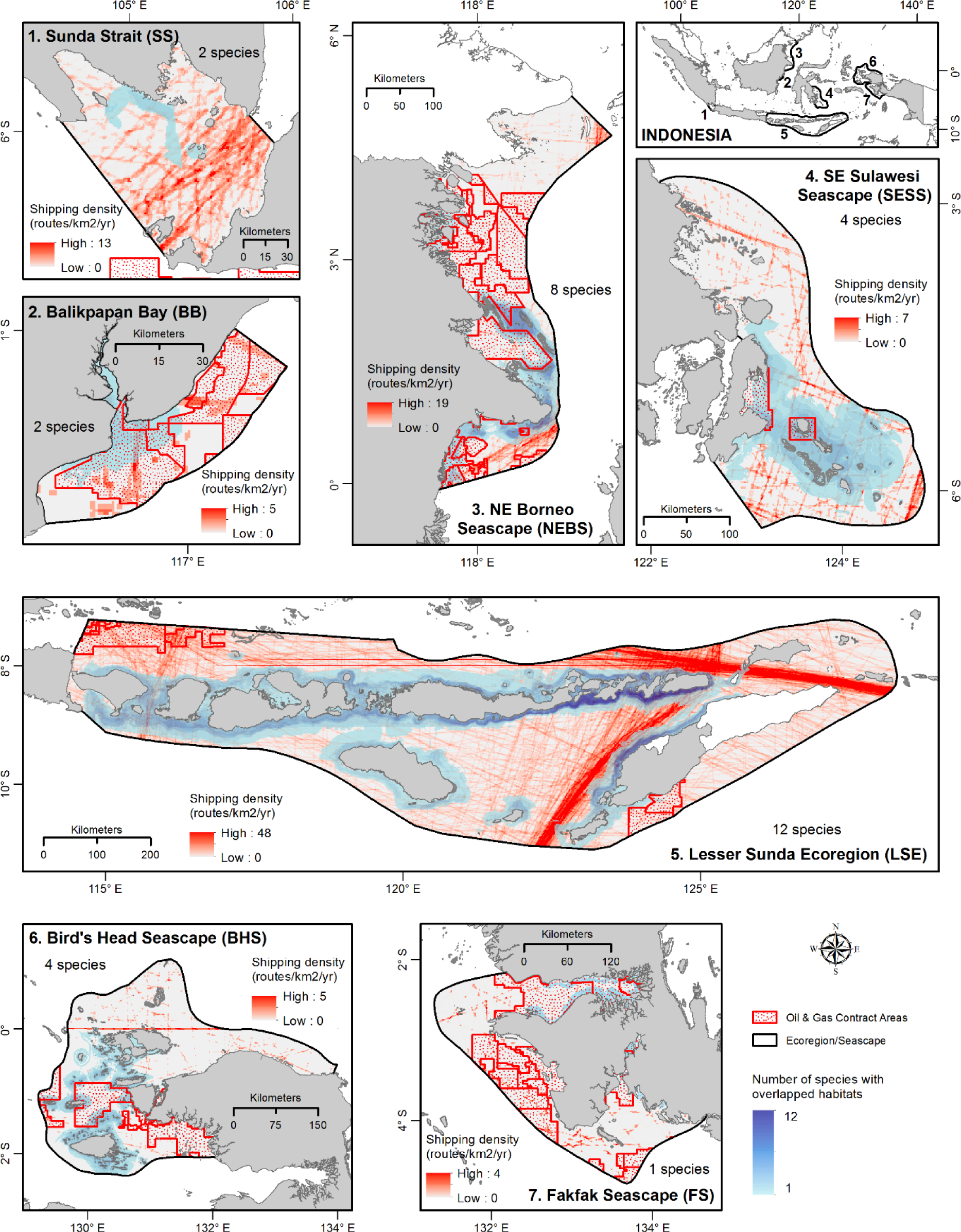
Species combined maps (the suitable habitats for at least one species, gradual blue colours showing number of species in each region) overlapped with two anthropogenic threats: marine traffic indicated by density of large vessel (>300 tons) shipping density (gradual red colours, data from Halpern et al., 2008), and oil & gas contract areas (red dotted areas). The highest number of shipping density and the number of overlapped species habitats are different in each region.

Superimposing the combined maps with MPAs and MSPs indicates areas of potential further protection, while superimposing with the anthropogenic threats indicates areas of high potential risk. Some highly suitable areas were outside protected areas where multiple human activities are allowed. Still, many of these suitable areas will potentially be covered in the MSP jurisdiction (e.g. in the SS, BB, NEBS, LSE and BHS, Figure 3), therefore it is important to take these critical habitats into account during MSP designation. Some suitable habitats are already included in MPAs (Figure 3), while some other areas coincide with oil and gas exploration areas and marine traffic (Figure 4). The majority of areas with predicted high habitat suitability were with low risk as can be seen for shipping traffic in the BB, NEBS, SESS, BHS, and FS (Figure 4). The LSE and SESS have large areas with suitable habitats overlapping with MPAs (Figure 3), and less overlapping with oil and gas concession areas (Figure 4). The LSE, however, and the SS have more dense shipping lanes suggesting that cetaceans face relatively substantial risk in both regions. Interestingly, several MPAs also overlap with oil & gas concession areas in five regions: the NEBS, SESS, LSE, BHS, and FS. The rest of the overlapping areas are considered to have low exposure to the two anthropogenic threats.

## 4. Discussion

The current knowledge of cetacean spatial distribution, its protection coverage and anthropogenic threats in Indonesia is still very limited though crucial for effective conservation management. Here we present the distribution modelling of 15 species in seven ecoregions or seascapes in Indonesia at a 1 km^2^ fine spatial resolution. The results reveal partial overlap with current MSP and MPA management systems and conflicts with anthropogenic activities. This study is a first attempt to provide more comprehensive spatial information using a currently available large collection of cetacean data. We summarize the main findings and general patterns, highlight remarkable results and provide the full information in the supplementary. Below we discuss (i) the spatial distribution resulting from the models, (ii) the quality of the models and their limitations, and (iii) the combined map and overlapping areas, as well as the potential uses of the model outputs, its implications for national cetacean conservation management and future perspectives.

### 4.1. Spatial distribution and habitat suitability predictions

Great heterogeneity in spatial distribution and habitat preferences among species and within species among different regions was identified from our work (Figure 2). This is not surprising taking into account the different ecologies of the 15 species. For cetaceans, utilizing spatially different habitats may be a strategy to minimise interspecific food competition (Bombosch et al., 2014). The heterogeneity reflects an interrelated influence of both topographic and oceanographic variables on the distribution of cetacean species. Both variables play a major role in partitioning the distribution of cetacean species, underlining how these variables were employed in the Maxent model to predict species distribution (Azzellino et al., 2012). The predictions reflect species’ traits in how they use the habitat. The fine scale of the present study allows us to identify habitat use and preferences in relation to different environmental features.

The species-specific habitat preferences of cetaceans are reflected in the set of environmental variables selected by each species’ model output. Both the composition and the number of environmental variables that determine habitat preferences, as well as the relative importance of the respective variables, are species-specific (Table 3). In most regions, each species mainly had 2 important variables determining its habitat that thus were retained in its model. Only *S. longirostris* in the SESS had 4 variables, the highest number of variables, retained in the model compared to other species model outputs (Table 3). Four other species in the LSE and 2 species in the NEBS also had 3 important variables. The high number of variables retained in the models indicates that their environmental niche in the region depends on a combination of several conditions. Despite the different important variables across the model outputs for each species and region, in general, Chl and SST (oceanographic predictors), and bathymetry, distance to coast and distance to −200m isobath (topographic predictors), were the most important variables that contributed to the distribution of most species in many regions. To a lesser extent SSS and distance to −1000m isobath were important for estuarine and oceanic species respectively. The ecological interpretation of complex relationships detected through Maxent model, of course, needs further analysis and validation.

#### Chlorophyll-a as a proxy of high productivity areas

Areas with high predictive suitability in every region were mainly related to high coastal or insular-reef complexity. In general, suitable habitat seems chiefly associated with high productivity, often with upwelling-modified waters close to topographic features (Redfern et al., 2017), which would create favourable foraging conditions. It has been reported before that cetacean distributions and population densities reflect the oceanographical conditions that are associated with biological productivity and diversity hotspots (Cama et al., 2012; Scales et al., 2014; Tobeña et al., 2016; Worm et al., 2005). That is why most suitable habitat areas are identified in our results mainly by chlorophyll-a concentration (Chl) and sea surface temperature (SST), and for a few species by sea surface salinity (SSS) (Figure 2, Table 3). Chl indicates phytoplankton and is the basis of the food web and is driven by nutrient availability (for example from rivers, and thus related to lower SSS) and higher SST (La Manna et al., 2016). Therefore it works as a good proxy for other bio-ecological factors (Moura et al., 2012) such as the distribution of zooplankton feeding on the phytoplankton on which the next predators feed including the prey species for the cetaceans. Thus, via this indirect link between primary biomass as represented by Chl and cetacean occurrence, Chl seems useful in identifying hotspots where cetaceans may aggregate. Our results are in accordance with Cotté et al. (2010) who reported that densities of dolphins were related to high Chl and high gradients of SST. All study areas are situated in the Indonesian tropical upwelling system, where oceanic currents strongly influence the temperature and primary productivity (Drushka et al., 2010; Steinke et al., 2014), benefitting marine species at all levels of the food web, including cetaceans. The distribution of some species (e.g. *S. sahulensis* in the FS and three species in the BHS), however, seemed not to be related to primary production but was instead strongly influenced by topographic variables (mainly distance to coast and bathymetry).

#### Topographic complexity indicating high habitat and prey diversity

Our model outputs clearly show that most species occurred in more complex topographic areas such as complex reefs and oceanic insular areas identifiable by certain depth e.g. −200m and −1000m isobaths, and in a few species by slope. In the NEBS and SESS, for instance, sperm whale distribution is closely associated with the −200m and −1000m isobaths. Distance to −2500m isobath, however, was not retained in any model output as an important variable, indicating that this variable is weak in determining cetacean distribution. Topographic variables have been suggested to influence and even drive persistent hydrographic features which can lead to the creation of predator hotspots (Bouchet et al., 2015; Pirotta et al., 2011; Praca et al., 2009) and most likely affects the availability, distribution and concentration of prey species (Arcangeli et al., 2016; Naud et al., 2003). Areas of high topographic heterogeneity and sea currents result in formation or localised upwelling, stimulating primary productivity that can sustain a rich food web structure and contain dense patches of prey, attracting large predators (Breen et al., 2016), including cetaceans. Blue whale in the LSE was predicted to be distributed in deep yet near coast areas around the strait between Flores and Sumba islands (Figure 2q). This is in agreement with what Ilangakoon and Sathasivam (2012) suggested that blue whales can be present in relatively small, localized highly productive feeding areas associated with strong upwelling year-round in Sri Lanka, and only make localized movements within this area. *O. brevirostris* prefers shallow, sheltered estuaries, such as in the NEBS, since such areas are commonly highly productive systems that can attract fish and top predators (Passadore et al., 2018). It has been shown before that distinct isobaths (−200m and −1000m) are important factors in determining the distribution of many cetaceans known to forage on pelagic schooling fish or deep-water prey (Goetz et al., 2015; Scales et al., 2014). In our study, especially the LSE, NEBS and SESS have very steep slopes with bathymetry exceeding −1000m within a short distance from shore. Deep yet near coast waters with complex topographic features are supportive of high cetacean diversity.

#### Species traits reflected in their predicted distribution

Cetaceans greatly differ in traits such as food choice, body surface-volume ratio and physiological adaptations for deep diving to forage (Mannocci et al., 2014). For instance, *P. macrocephalus* and *G. melas* have lower energetic costs and are able to forage deeper than most delphinid species and therefore can exploit deeper food resources along the continental slope than species with shallower diving capabilities (Mannocci et al., 2014). Predicted habitat from our study was also higher for most species in coastal (mainland or insular) areas. Cetaceans increase their benefits by spending as much time as possible close to the areas where the likelihood of finding their preferential prey may be higher (La Manna et al., 2016). For instance, bottlenose dolphins in all regions prefer coastal areas which have shallower feeding grounds that often host complex and rich food webs (La Manna et al., 2016), which is optimal for a species that chiefly forages on demersal prey (Bearzi et al., 2008).

The same species can have distinct habitat preferences in different regions and a combination of different environmental variables may influence its predicted distribution. The expression of the combination of variables can be unique in each ecosystem (Redfern et al., 2017). Our results showing that sperm whale can inhabit different areas with variable characteristics are in agreement with those reported by (Praca et al., 2009), since some cetaceans tend to be opportunistic towards their surrounding available habitats. In this case, sperm whale can sustain their needs in both productive coastal and complex insular-reef areas, as well as in more offshore oligotrophic areas (Lambert et al., 2014). Comparison of the habitat preference of a species over different regions characterized by different complex environmental variables could add understanding on the core qualities of the species’ favourable areas. Habitats overlapping for different species are especially important to study further as understanding co-occurrence due to niche specialisation as well as potential interspecies competition better might be very helpful for cetacean conservation management efforts.

### 4.2. Quality and limitations of the models

To be able to build a plausible habitat suitability model we carefully implemented data, performed predictor quality control (e.g. reducing data autocorrelation and environmental variable multicollinearity) (Brown et al., 2017), chose Maxent as it is specifically designed to handle presence-only data (Elith et al., 2006; Phillips et al., 2006), and corrected over-prediction model outputs (Brown et al., 2017). Model evaluation metrics (based on moderate-high AUC and low SD values) indicate that most of model outputs had internal model consistency, thus supporting the reliability of models, and the distribution models produced had reasonable robustness (Bombosch et al., 2014). The results are useful not only for ecological investigation but could also support conservation management decision making. Nevertheless, in the interpretation of model outputs attention should be paid to some methodological limitations. Some common issues derived from sample size, different sources of data, sampling bias and different spatial scales could influence the accuracy of the Maxent algorithms (Elith et al., 2006) and are discussed below.

The sighting data used in this study was collected opportunistically in areas of known occurrence or areas that are easy to survey; hence our sighting data represents a sampling bias (Figure 1). Spatial autocorrelation, the tendency of locations close to one another to share similar values for environmental variables, is also common in species occurrence data (Dormann et al., 2007). This can lead to a number of problems in species distribution modelling, including biased coefficient estimates, inflated measures of model evaluation and difficulties in transferring predictions in geographical space (Guélat and Kéry, 2018). Spatial autocorrelation arises from several processes, but mainly by sampling bias (El-Gabbas and Dormann, 2018). To correct for non-homogeneous distribution of the sampling effort (Phillips et al., 2009) we applied the ‘bias file’ function in the Maxent models (Brown, 2014; Brown et al., 2017).

Due to sample size limitations, we only modelled the habitat suitability for 15 out of 34 cetacean species in seven regions in Indonesia (Figure 2, Table S1). Obtaining enough data for modelling the habitats of cetaceans, or top predators in general, is challenging because these organisms are by nature sparsely distributed compared to lower trophic levels, and their detection is often imperfect, often resulting in scarce datasets (Virgili et al., 2017). We performed the distribution models, despite the relatively small number of sightings for some species, as a useful first approach of cetacean habitat modelling that could accelerate the knowledge on distribution of these never-studied populations. A number of previous studies also successfully used distribution models for multiple cetacean species with low numbers of sighting records (e.g. Breen et al., 2016; Correia et al., 2015). For some species in our study the number of sightings unfortunately was too low (between 10 and 15 points) for our distribution modelling, highlighting the need for further dedicated monitoring of cetacean presence in this area.

Among available presence-only models, we chose Maxent because it appeared more suitable to model the predictions of species distribution with complex interactions between the response and the predictor variables (Elith et al., 2011; Phillips and Dudík, 2008) and seemed to manage datasets characterised by scarce data well (Wisz et al., 2008). The use of Maxent, along with limited occurrence data, can be a cost-efficient way to obtain information for regions where no preceding studies exist. Ideally, the number of occurrence data point is >15 sightings to prevent inconsistent model results (Aguirre-Gutiérrez et al., 2013), although Maxent is also known to perform well with small sample sizes (Pearson et al., 2007), which was the case for several species in our study. With more sightings over the year it also becomes possible to generate seasonal predicted distributions for species, which is especially relevant for non-resident species. Despite the aforementioned limitations, our results provide crucial first insights to support conservation and management strategies where there was previously a fundamental lack of knowledge regarding cetacean distribution.

### 4.3. Overlapping suitable habitats and conservation management areas and threats

This study highlights the importance of several ecoregions and seascapes in Indonesia for cetacean species, especially in the LSE and NEBS, while several species could not even be included due to insufficient data. The more species that occur in a region, the more important the region becomes from a conservation perspective (Zacharias and Gregr, 2005). Most of the important coastal habitat areas also contain major human settlements, harbours and tourism destinations. These anthropogenic activities pose actual and potential future threats to cetacean populations in those regions. The combined species maps also show that important suitable cetacean habitat is in the vicinity of insular areas. Understanding the current habitat use of cetaceans is a necessary step in comprehending and evaluating the effects of human activities on the species to be followed by conservation management strategies including MSP and MPA development. In spatially explicit risk assessments the local species distribution should be linked to the potential effects and distribution of human activities (Stelzenmüller et al., 2009), and eventually measures can be developed to reduce or avoid the adverse impacts. This study provides cetacean distribution maps at a fine (1 km^2^) resolution which is useful for strategic MSP of human activities that are potentially harmful to cetaceans (Roberts et al., 2016).

Using the Maxent model to identify cetacean suitable habitat can provide the scientific justification for their more effective protection. Our results show that large areas of important cetacean habitats are currently not protected in the existing conservation management system, since areas of highest probability were located outside MPAs (Figure 3). These areas could be considered in other site-based protection approaches such as MSP development (Coomber et al., 2016). The information on spatial gaps in protection coverage is useful to assess effectiveness of existing MPAs for conserving biodiversity in the region. To our knowledge, no studies so far have investigated whether Indonesian MPAs are adequately protecting marine mammals, which is the ambition of the Indonesian government (Sahri et al., 2020). The different extent of species habitats makes the delineation of MPAs a challenge. The most suitable cetacean habitats in coastal and complex insular-reef areas deserve special protection against human activities that may threaten cetaceans, and deserve special attention when establishing MPAs. Typically small MPAs offer limited conservation benefits (Gaines et al., 2010) particularly for mobile species. Therefore the size and design of MPAs and consequent management measures should be adapted to the ecological principles of the cetacean life cycles and that of their prey (Mangano et al., 2015). MPAs also need to be viewed in the larger context of the entire ecosystem including the extent to which these habitats interact at larger spatial scales (Azzellino et al., 2012). In Indonesia, provincial MSP jurisdiction only covers waters until 12 nm from the coastline. Thus, to improve the protection of cetaceans outside current MPAs and provincial MSPs, the suitable habitats should be taken into account in open-sea MSP establishment (The Government of The Republic of Indonesia, 2014b). The existing knowledge on cetacean habitats in this region, fortunately, has been largely included as Important Marine Mammal Areas (IMMA) (IUCN-MMPATF, 2019). In addition, the ecosystem in which marine mammals live often encompasses the waters of more than one country (Giannoulaki et al., 2017). This is the case in the LSE, where two countries (Indonesia and Timor Leste) share the coastline. Our results allow the identification of the suitable habitat for the species over wider areas, providing the means to strengthen protection for the species beyond an individual country’s territorial waters.

Worldwide, oil-gas exploration is currently expanding into areas previously undisturbed by industrial development (Wilson et al., 2013), including areas important for cetaceans. The BB and NEBS in East Kalimantan, and the FS in Papua hold significant hydrocarbon reserves and are well-known as important national oilfields (Patra Nusa Data, 2016) that can support the national demand for oil/gas and income. The areas overlap with important cetacean habitat identified from our study. The concession areas are subject to seismic activity for hydrocarbon deposits in the seabed during exploration activities. Cetacean mass stranding events have been related to seismic exploration (Filadelfo et al., 2009; Frantzis, 2004). Seismic survey is also one major contributor of anthropogenic ocean noise (Elliott et al., 2019; Farmer et al., 2018) that cetaceans are exposed to, in addition to the noise produced by marine shipping. Although some cetacean suitable habitats overlap with oil-gas concession areas, Indonesia has no regulation on underwater noise such as from seismic activities (Sahri et al., 2020). Oil spills from exploration and production activity have been reported to cause massive pollution to the marine environment (Eckle et al., 2012; Farmer et al., 2018). Cetaceans suffer from oil spills by direct contact with crude-oil or high concentrations of volatile gases, by indirect exposure to toxic oil hydrocarbons via their prey or loss of prey, as well as from oil spill response activities including increased vessel operations, dispersant applications, and oil burns (Dias et al., 2017; Schwacke et al., 2017). The cetacean habitat maps from this study identify potential overlapping areas of conflict, which could be avoided. Where potential threats are already well known, such as near production wells for oil and gas extraction, enhanced preparedness for spill events in proximity to species habitat is also worthy.

Shipping is one of the world’s largest industries and dominates ocean use (UNCTAD, 2017), thus excluding this sector from decision-making could lead to increased conflict among user groups and negative environmental impacts (Coomber et al., 2016; Metcalfe et al., 2018). This marine transport may directly affect the habitat use of marine mammals, since they coincide in the overlapping ocean spaces, and if uncontrolled, can create adverse impact to the species through collision and noise pollution. The overlapping maps of suitable habitat and shipping density from our study showed two regions, the SS and LSE, had high exposure to marine traffic (Figure 4), therefore mitigation for the anthropogenic threats should be focused on these areas. Within Indonesian waters, three archipelagic sea lanes (Indonesian: *Alur Laut Kepulauan Indonesia*, ALKI) have been established i.e. in Sunda Strait (ALKI I), Lombok Strait (ALKI II), and Ombai Strait (ALKI III). ALKIs are some of the busiest shipping routes in the Asia and Oceania regions, with an estimated 27% of the world’s oil traffic per day being transported through the Malacca Straits and the ALKIs (Asian Development Bank, 2014). Although some cetacean suitable habitats exist close to the designated ALKIs, Indonesia has not declared any sensitive areas to protect such habitats from possible ship collision, noise and oil pollution produced by shipping operations. The current shipping is not only concentrated in the established lanes, but also occurs in unofficial shipping lanes northeast of the ALKI III (Figure 4). These lanes are also important migratory corridors for a variety of marine mammals including pygmy blue whale, as shown by telemetry data (Double et al., 2014). During 1975– 1997, oil spills from 104 shipping accidents polluted Indonesian marine and coastal areas (Nontji, 2000). A better understanding of the shipping intensity overlapping with cetacean habitat can be used to inform ecosystem-based management within an MSP framework (Coomber et al., 2016). For instance, the information can be useful in planning low-impact shipping corridors, informing the delineation, modification or adjustments of the shipping lanes and additional protection needed for cetaceans by establishing new marine reserves (Dawson et al., 2018; Dransfield et al., 2014).

It is important to note that the spatial footprint of marine traffic used in this study was only based on large vessels, so is an underestimation of the shipping intensity. As mandated by the International Maritime Organization (IMO), only large vessels (>300 tons) making international voyages and all passenger ships regardless of their size are required to have an Automatic Identification System (AIS) to track their movements in real time (Shelmerdine, 2015). Small recreational boats do not have AIS transmitters and can exist unnoticed in large numbers in certain areas. Vessel Monitoring System (VMS) data from fishing vessels unfortunately is not yet publicly available in Indonesia. Therefore, our marine traffic information excludes a large portion of smaller vessels, many of which travel a lot. Furthermore, these smaller vessels tend to operate inshore which will particularly affect exposure for the cetaceans which have been shown to prefer shallower coastal waters. Further work on improving our knowledge of the spatial distribution of inshore marine traffic is imperative and will give new insights into possible boat collision and noise risk.

### 4.4. Applications for national cetacean conservation and management, future perspectives

The risk exposure maps (Figures 3 and 4) from our study can support management decisions and conservation measures as in MSPs, although they are only one input into a systematic conservation planning process (Margules and Pressey, 2000). Our analysis only included two anthropogenic threats: marine shipping and oil and gas industry. Fisheries, tourism and many more human activities are synergistically or individually also important anthropogenic threats to cetaceans (Thorne et al., 2012; Wise et al., 2019). More accessible information is needed to better understand the actual allocation of these other anthropogenic threats and the overlap with cetacean distribution, MPAs and MSP. The present study is an important step as a basis for assessing the overlap between cetacean habitat, anthropogenic threats, MPAs and MSP. The cetacean spatial prediction maps as presented here provide a basis for a valuable planning tool in the context of anthropogenic threats, and can also direct additional monitoring efforts to further improve the spatial and species coverage. The inclusion of other anthropogenic pressures is a next step to better understand how human activities overlap with cetacean distribution and to determine where MSP for conservation management is urgently needed.

## 5. Conclusion

The cetacean distribution maps presented here provide a recent habitat preference characterization of the studied species. Closer analysis clearly indicates why Indonesian waters are rich in cetacean diversity, as topographic and oceanographic factors favour primary production and habitat complexity including deep waters. We provide fine resolution distribution maps of 15 cetacean species in seven regions in Indonesia. We highlight the areas of highly suitable cetacean habitat as priority areas for future spatial conservation decisions, for example, in the SS, the BB, the northern part of the NEBS, and the north-eastern part of the LSE. Our results also identified potential areas of conflict with human activities. High risk areas will vary regarding seasonal distribution; further research on seasonal modelling is then needed e.g. to find the best time for performing oil and gas exploration (seismic activity). It is important to realise that 19 out of the 34 species reported were not included in this modelling study because of lack of sighting data. The distribution of these species should also be taken into account in future cetacean conservation management planning.

The spatial planning efforts should also include a larger spatial scale than 12 nm to ensure all important biodiversity assets are appropriately represented in new MPAs, in protected zones within provincial MSP jurisdiction, or in future open-sea MSP establishment beyond 12 nm. Given current global environmental changes, dedicated monitoring of possible changes in cetacean distribution is necessary for adaptive conservation management. Our species distribution prediction maps can help direct future monitoring surveys to collect more information on understudied species as well as of predicted suitable habitat for which no observations are available yet.

The approach we present is relatively inexpensive, but parallels and is complementary to large-scale marine mammal surveys, providing a model that can be used where resources are limited. This approach can be applied anywhere to model cetacean habitat use, identify priority areas for conservation, and highlight potential areas of conflict with human activities to inform conservation management.

## Acknowledgements

The authors would like to thank the people and institutions that provided data and contributed to this work: Yusuf Fajariyanto, Purwanto, Awaludinnoer Ahmad and The Nature Conservancy Indonesia, Sugiyanto and WWF Indonesia, Wakatobi National Park Authority and all staff who mainly collected data in the field, Yayasan Konservasi RASI and all volunteers, Hadi Yoga Dewanto and the Ministry of Marine Affairs and Fisheries of Indonesia for MPA data, Rokhman Khabibi for the oil and gas contract area map, and Doddy Yuwono and the Geospatial Information Agency of Indonesia for coral reef GIS data. Some data was obtained through data sharing agreements, and is therefore not publicly accessible. This work was financially supported by the Indonesia Endowment Fund for Education (LPDP) Scholarship from the Ministry of Finance of Indonesia for the first author [contract number: PRJ-482/LPDP.3/2017].

